# Phthalates Impair Estrogenic Regulation of HIF2α and Extracellular Vesicle Secretion by Human Endometrial Stromal Cells

**DOI:** 10.1101/2024.12.12.628185

**Authors:** Jacob R. Beal, Arpita Bhurke, Kathryn E. Carlson, John Katzenellenbogen, Jie Yu, Jodi Flaws, Indrani C. Bagchi, Milan K. Bagchi

## Abstract

Di(2-ethylhexyl) phthalate (DEHP), a known endocrine-disrupting chemical, is a plasticizer found in many common consumer products. High levels of DEHP exposure have been linked to adverse pregnancy outcomes, yet little is known about how it affects human uterine functions. We previously reported that the estrogen-regulated transcription factor hypoxia-inducible factor 2 alpha (HIF2α) promotes the expression of Rab27b, which controls the trafficking and secretion of extracellular vesicles (EVs). EVs facilitate communication between multiple cell types within the pregnant uterus, ensuring reproductive success. In this study, we report that exposure of differentiating primary human endometrial stromal cells (HESC) to an environmentally relevant concentration (1 μg/mL) of DEHP or its primary metabolite mono(2-ethylhexyl) phthalate (MEHP) markedly reduces the expression of *HIF2α*. We also observed a concomitant decrease in *RAB27B* expression, reducing EV secretion from HESC. Interestingly, we found that DEHP or MEHP exposure disrupts estrogenic regulation of the HIF2α/Rab27b signaling pathway.

Estrogen receptor alpha (ERα) could no longer bind to the *HIF2α* regulatory region following phthalate treatment, and epigenetic analysis suggested that this may be due to hypermethylation of nearby CpG islands. Further investigation revealed a potential interaction between ERα and the transcription factor Sp1 within the HIF2α regulatory region, which is affected by the inhibition of Sp1 binding to the phthalate-induced hypermethylated DNA. Additionally, our results suggest that the abnormal DNA methylation is likely due to increased expression of the DNA methyltransferase 1 (*DNMT1*) gene in response to phthalate exposure. Overall, this study provides valuable mechanistic insights into how phthalate-induced differential DNA methylation disrupts estrogenic regulation of the *HIF2α* gene and, consequently, EV secretion during HESC differentiation. This knowledge is crucial for our understanding of how phthalates may cause adverse reproductive outcomes by disrupting the hormonal regulation of cell-to-cell communication within the pregnant uterus.

## Introduction

Di(2-ethylhexyl) phthalate (DEHP) is a member of a class of compounds called phthalates, which are synthetic chemicals that are used as plasticizers (1). DEHP can be found in plastic products such as packaging, toys, medical devices, building materials, and cosmetics. Since phthalates do not covalently bind to plastic polymers, they can leach out of these products leading to human exposure. Humans can be exposed to phthalates via multiple routes, such as ingestion, inhalation, and dermal contact. Environmental exposure to phthalates is so prevalent that nearly 100% of tested urine samples contain phthalate metabolites (2–4).

DEHP is one of the most commonly found phthalates in the environment, with daily human exposure to DEHP. having been estimated to be around 3-30 μg/kg/day. Some studies report a higher daily intake level in women compared to men (5–7). Unfortunately, DEHP is a known endocrine-disrupting chemical (EDC) that negatively affects the female reproductive system, as well as other aspects of human physiology. DEHP exposure *in utero* has been associated with shorter pregnancy duration (8), increased incidences of miscarriages (9), and delayed puberty in girls (10).

*In vivo* studies in rodents have confirmed the contribution of DEHP in disorders of the female reproductive tract. Previous studies have found that chronic exposure to DEHP at various doses affects the estrous cyclicity of female mice (11,12). DEHP exposure has been reported to adversely impact uterine functions with certain doses causing a reduction in epithelial cell proliferation and an increase in abnormally dilated blood vessels in the endometrium (13).

Concurrent DEHP exposure and a high-fat diet have also been demonstrated to lead to abnormal placentation in mice (14). Chronic exposure to phthalates in mice has been reported to alter pubertal onset, affect folliculogenesis, and decrease fertility (15–17). Some of the effects have even been shown to be transgenerational, with subsequent non-dosed generations exhibiting early pubertal onset, altered estrous cyclicity, and reduced fertility (17). Transgenerational toxicity of DEHP exposure has been well documented and inherited dysregulated epigenetic markers of gene expression has been posited by multiple groups as a mechanism of transgenerational toxicity (18–20).

During early pregnancy, events critical to reproductive success, such as implantation, decidualization, maternal angiogenesis, and placentation, involve multiple tissues within the uterus and as such must be intricately coordinated to properly support embryonic development (21). Previous work from our lab described how the maternal endometrium facilitates communication between different cell types to coordinate these critical events. We reported that, in pregnant mice, endometrial stromal cells, while undergoing differentiation to secretory decidual cells in a process termed decidualization, induced the expression of transcription factor hypoxia-inducible factor 2 alpha (*Hif2α*) in response to estrogenic signaling (22). This is not surprising, as a hypoxic condition is known to prevail in the uterus of many species during early pregnancy (23,24). The adaptation to hypoxia, mediated by HIF2α, is critical as genomic deletion of the *Hif2a* gene in mouse uterus leads to implantation failure and female infertility (22).

HIF2α was shown to promote the expression of vesicular trafficking protein Rab27b (22). Later work demonstrated that Rab27b controls the secretion of extracellular vesicles (EVs), which were found to mediate functional communications between different cell types located in the uterus via transfer of key protein cargoes enclosed within the EVs (25). The HIF2α/Rab27b/EV secretion pathway was also found to be conserved within primary human endometrial stromal cells (HESCs) and EVs derived from differentiating HESCs were shown to promote decidualization, angiogenesis, and trophoblast differentiation (26).

Estrogenic regulation of *HIF2α* and the EV signaling pathway that HIF2α governs is an important mediator of cell-cell communication within the uterus during early pregnancy. If we find that this endocrine signaling pathway is perturbed by exposure to phthalates, it will enhance our knowledge of how phthalates act within the uterus, bolstering our understanding of how phthalates may cause adverse reproductive outcomes. Therefore, we conducted a study to test whether DEHP disrupts the HIF2α/Rab27b/EV secretion pathway that mediates cell-cell communication during early pregnancy. Since the signaling pathway is conserved between the mouse and the human, we elected to treat primary HESCs cultured in hypoxia under decidualization conditions with DEHP to determine if, and how, phthalate exposure disrupts this process. Since, in the body, DEHP is rapidly metabolized, we also treated the cells with DEHP’s primary metabolite, mono(2-ethylhexyl) phthalate (MEHP) (27), to compare its effects to its parent compound. We performed gene expression and epigenetic analyses, quantified EV secretion, and measured transcription factor DNA occupancy to illustrate the impact of phthalates on this hypoxia-induced cell signaling pathway crucial during early pregnancy.

## Methods

### Primary HESC Culture

Deidentified primary HESCs were collected following the guidelines for the protection of human subjects participating in clinical research approved by the Institutional Review Board of Wake Forest School of Medicine. Samples were collected from fertile women during the proliferative stage of the menstrual cycle by biopsy under anesthesia before laparoscopy as described previously (28). Donors ranging in age from 28 to 42 years in parity from 1 to 2 provided written informed consent and were confirmed to display no signs of endometrial pathologies. HESCs were isolated as previously described (28).

HESCs were passaged under normoxic conditions (20% O2) while cultured in Phenol red-free Dulbecco’s modified Eagle medium (DMEM)/F-12 (Gibco) supplemented with 5% charcoal dextran-stripped FBS (Atlanta Biologicals), 50 μg/mL penicillin, and 50 μg/mL streptomycin (Invitrogen). For *in vitro* differentiation experiments, HESCS were moved to a hypoxia incubator (3% O2) and treated with a differentiation cocktail containing 1 μM progesterone (Sigma-Aldrich), 10 nM 17-β-estradiol (Sigma-Aldrich), and 0.5 mM 8-bromo-adenosine-3’,5’-cyclic monophosphate (Sigma-Aldrich) in Phenol-red free DMEM/F-12 medium (Gibco) supplemented with 2% exosome-depleted FBS (Gibco). In some experiments, dimethyl sulfoxide (DMSO) (Sigma-Aldrich), 0.1 or 1 μg/mL DEHP (Sigma-Aldrich), or 1 μg/mL MEHP (AccuStandard) were added to the differentiation cocktail. DMSO was used as a vehicle to create stock solutions of DEHP and MEHP. Differentiation was allowed to take place over 72 hours before sample collection. DC and DEHP or MEHP treatment (if applicable) was replenished after 48 hours.

### siRNA-mediated knockdown

Primary HESCs were transfected by siRNA targeting *ESR1* or *SP1* gene or scrambled siRNA (Dharmacon) following the manufacturer’s protocol (siLentFect; Bio-Rad). Briefly, a final concentration of 20 nM siRNA was mixed with siLentFect lipid reagent to transfect cells. Cells were incubated with siRNA-lipid mixture overnight and then the media was replaced with media containing DC to induce differentiation as described above.

### RNA Isolation and qPCR analysis

After 72 hours of differentiation, total RNA was extracted from HESCs using TRIzol (Invitrogen) following the manufacturer’s instructions. RNA was converted to cDNA using AffinityScript Multiple Temperature Reverse Transcriptase kit (Agilent) according to the manufacturer’s instructions. Quantitative PCR analysis was performed on the cDNA using Power Sybr Green PCR master mix (Applied Biosystems) and gene-specific primers (IDT). *36B4* was used as a housekeeping gene. For each treatment condition the mean cycle threshold (Ct) was calculated from three replicates of each sample. ΔCt was calculated as the mean Ct of the gene of interest subtracted by the mean Ct of the housekeeping gene (*36B4*). ΔΔCt was calculated as the difference of ΔCt of experimental and controls. Fold change of gene expression relative to control was computed as 2^-ΔΔCt^. The data was then expressed as mean fold induction and standard error of the mean (SEM) calculated from 2-6 independent experiments.

### EV Isolation and Quantitation

Conditioned media were collected after 72 hours from cells undergoing DC and phthalate or siRNA treatment as described above. Conditioned medium was centrifuged at 3,000 g for 10 minutes to obtain a cell-free specimen. The supernatant was used to extract EVs using miRCURY Exosome Cell/Urine/CSF (Qiagen) following the manufacturer’s protocol. The harvested EV pellet was resuspended in 0.2 μm filtered PBS and a sample was taken for quantification by microfluidic resistive pulse sensing (MRPS).

MRPS was performed using a Spectradyne nCS1 instrument (Spectradyne) according to the manufacturer’s instructions. EVs were diluted in PBS + 1% Tween20 and EV suspension was loaded on polydimethylsiloxane cartridges. These factory-calibrated cartridges quantify particles only within specific size ranges. To quantify EV concentration, we utilized the C-400 cartridge to quantify particles from 65-400 nm in size as described previously (25,26). Measurements were given as a particle concentration (particles per mL) and data was presented as the average fold change in particle concentration compared to control and SEM calculated from 2-3 independent experiments.

### Chromatin Immunoprecipitation

One million HESCs that underwent DC (and in some cases phthalate treatment) as described above were collected after 72 hours treatment in hypoxia and used per reaction. ChIP experiments were carried out employing the ab500 ChIP Kit (Abcam) according to the manufacturer’s instructions. For the immunoprecipitation step, fragmented DNA was incubated overnight with 5 μg of anti-ERα (Abcam Cat# ab32063, RRID: AB_732249), or anti-HIF2α (Novus Cat# NB100-122, RRID: AB_10002593), or anti-Sp1 (Abcam Cat# ab231778, RRID: AB_2936949) or Rabbit IgG (Abcam Cat# ab172730, RRID: AB_2687931). After immunoprecipitation and DNA purification steps, DNA underwent qPCR. Primers were designed spanning putative estrogen response elements (EREs), hypoxia response elements (HREs), or putative SP1 binding motifs within the regulatory regions of *HIF2α* or *RAB27B*. Additional primers were designed at an ERE-, HRE-, and SP1 binding motif-free negative control locus.

Primer sequences are available upon request. Resultant Ct values were normalized to the negative control locus and data were shown as mean fold change relative to the IgG control and SEM computed from 3-4 independent experiments.

### Radiometric Competitive Binding Assay

Binding affinities of DEHP and MEHP were determined using a radiometric competitive binding assay, with tritiated estradiol used as a tracer and hydroxylapatite used to adsorb the ligand-receptor complex, as described previously (29). Human ERα was expressed in baculovirus and purified (Thermo Fisher Scientific, Catalog # A15674). Data are shown as relative binding affinity (RBA) values. where the RBA of estradiol is 100. RBA values were calculated from 2-3 replicates per test compound.

### Quantification of DNA Methylation

After 72 hours of treatment with phthalates and DC as described above, DNA was isolated from HESCs using DNeasy Blood & Tissue Kit (Qiagen) according to the manufacturer’s instructions. DNA methylation was quantified using the OneStep qMethyl kit (Zymo Research) following the manufacturer’s protocol. Primers were designed flanking at least two methylation-sensitive restriction enzyme sites surrounding an ERE in *HIF2α* regulatory region. Methylated sites in the region were protected from digestion by restriction enzymes and thus were not amplified by qPCR. This allows for the percentage of methylation to be estimated using the ΔCt of the test group (with restriction enzymes) and reference group (lacking restriction enzymes). Nearby CpG islands were identified using a computation method described previously (30) and primers were also designed within the two CpG islands identified. Representative data were then shown as mean fold change in % methylation compared to vehicle control and SEM computed from 2-3 independent experiments.

### Statistical Analysis

Statistical analysis was performed in all experiments using an unpaired Student’s t-test to determine if any treatment was statistically different from the control. GraphPad Prism Version 9 (GraphPad Inc.) was used for statistical analysis. A P-value of ≤ 0.05 was required to be considered statistically significant.

## Results

### Phthalate treatment impairs HIF2α regulated EV secretion in decidualized HESCs

During early pregnancy, endometrial stromal cells undergo an ovarian steroid hormone-regulated transformation where the stroma differentiates into a more secretory subtype termed the decidua. This differentiation process, known as decidualization, is marked by morphological and gene expression changes as well as an increase in EV secretion (25,26). Concurrently, the endometrium must adapt to a hypoxic environment, which it does by promoting the expression of transcription factor HIF2α.

To study these phenomena *in vitro*, we use a well-established system where decidualization is induced in HESCs in response to a differentiation cocktail (DC) consisting of 1 μM progesterone, 10 nM 17-β-estradiol, and 0.5 mM 8-bromo-adenosine-3’,5’-cyclic monophosphate. To test whether phthalate exposure disrupts this process, we initially added vehicle, 0.1 μg/mL, or 1 μg/mL DEHP to the DC. We chose these concentrations since DEHP concentration within the peritoneal fluid of healthy women is around 0.46 μg/mL (31), so adding 0.1 μg/mL or 1 μg/mL of DEHP in our experiments would be environmentally relevant concentrations to study phthalate effect.

After 72 hours of decidualization under hypoxic conditions, we observed that, compared to vehicle control, neither concentration of DEHP treatment triggered significant changes in the gene expression of traditional markers of human decidualization, insulin-like growth factor binding protein 1 (*IGFBP1*) or prolactin (*PRL*). However, 1 μg/mL, but not 0.1 μg/mL DEHP treatment caused a marked reduction in the expression of the *HIF2α* gene (also known as *EPAS1*) (**Fig. 1A**). Previous studies had shown that *HIF2α* gene expression is typically strongly induced in HESCs when cultured under decidualization conditions in hypoxia without treatment of phthalates (26).

**Fig. 1.**
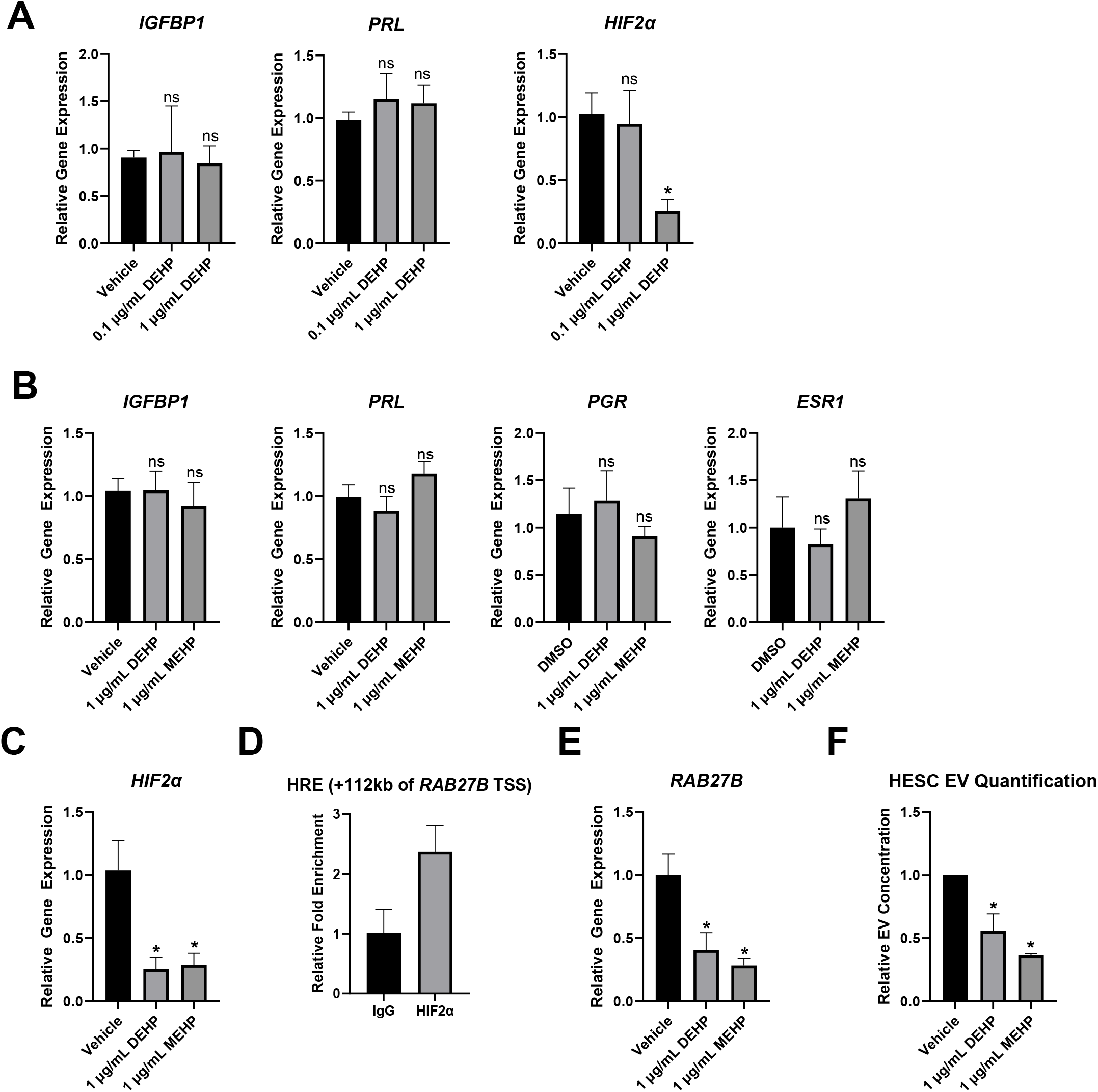
Treatment with DEHP or MEHP impairs HIF2α/Rab27b mediated EV secretion without affecting early HESC decidualization. **A**. HESCs were treated with a Decidualization Cocktail and vehicle, 0.1 μg/mL DEHP, or 1 μg/mL DEHP for 72 h under hypoxic conditions. RNA was extracted, and gene expression analysis was performed using primers specific for key decidualization markers, *IGFBP1 (left)* and *PRL (middle)*, as well as HIF2α (right). **B.** Gene expression analysis was also performed after treatment with 1 μg/mL DEHP or MEHP using primers specific for *IGFBP1 (left), PRL (middle left), PGR (middle right), and ESR1 (right). HIF2α* (**C**) and *RAB27B* (**E**) gene expression were also quantified. *36B4* was used to normalize gene expression. Data are shown as mean fold change ± SEM (n = 3-6 independent experiments), * p<0.05, relative to vehicle treatment. **D.** ChIP was performed on HESCs that had been treated with a Decidualization Cocktail for 72 h under hypoxic conditions. Chromatin enrichment was quantified by qPCR and normalized to a negative control locus. Primers flanking putative hypoxia response elements within the *RAB27b* enhancer region were used to quantify chromatin enrichment. Representative data are shown as mean fold change ± SEM (n = 2 independent experiments) **F.** EVs were isolated from conditioned media of an equal number of differentiating HESCs treated with vehicle, DEHP, or MEHP using the miRCURY (Qiagen) kit and analyzed by MRPS (Spectradyne, LLC). The concentration of EVs ranging from 60 to 400 nm was quantified. Data are shown as mean fold change ± SEM (n = 4 independent experiments), * p<0.05, relative to vehicle treatment.

Efficacy of the 1 μg/mL dose of DEHP in reduction of critical *HIF2α* gene expression prompted continued investigation into the response of HESCs to this environmentally relevant concentration of phthalate exposure. Further, we examined the effects of DEHP’s primary metabolite, MEHP, at this concentration to determine if the cellular response to phthalate exposure remained the same as its parent compound. We found that previously mentioned decidualization markers IGFBP1 and PRL remained unaffected by MEHP exposure. Similarly, gene expression of significant markers and regulators of HESC decidualization, progesterone receptor (*PGR*) and estrogen receptor alpha (*ESR1*), remained unchanged after exposure to either DEHP or MEHP (**Fig. 1B**). MEHP exposure also exhibited a similar response in *HIF2α* gene expression (**Fig. 1 C**).

Our next experiments determined if phthalate exposure interfered with the adaptive signaling pathway that HIF2α mediates in differentiating HESCs in response to hypoxia. HIF2α has been shown to regulate *RAB27B* expression (22,26), and Rab27b has been shown to facilitate EV secretion (26,32). We confirmed via ChIP-qPCR that HIF2α directly regulates *RAB27B* gene expression during HESC differentiation through binding at an enhancer site in its regulatory region (**Fig. 1D**). We then examined whether *RAB27B* gene expression was affected by exposure to phthalates. We found that *RAB27B* expression was indeed markedly reduced in response to DEHP or MEHP treatment (**Fig. 1E**). We also quantified the number of EVs secreted by HESCs during the decidualization process when exposed to phthalates. After 72 hours of differentiation in hypoxic conditions, while exposed to 1 μg/mL DEHP or MEHP or vehicle, we collected the conditioned media and isolated EVs using a miRCURY (Qiagen) kit. The EV pellet was resuspended in PBS, and its concentration was quantified by microfluidic resistive pulse sensing (MRPS). We observed a significant reduction in EV secretion after treatment with DEHP or MEHP (**Fig. 1F**), concomitant to the reduction in Rab27b expression in the same cells.

### Estrogenic regulation of EV secretion in HESCs is disrupted by phthalate treatment

To explore how phthalates may be affecting HIF2α-mediated EV secretion by HESC, we investigated the role of estrogen receptor signaling in this process. This investigation was prompted by existing literature that suggests a connection between estrogenic signaling, HIF2α gene regulation, and EV secretion. *Hif2α* expression was found to be induced by estrogen in pregnant mice (22), and a recent study described how 17β-estradiol increased the number of EVs secreted in ER+ breast cancer cells but not in ER-cell lines (33).

Therefore, we performed experiments to first determine if perturbation of ERα signaling affects the HIF2α/Rab27b/EV secretion pathway in differentiating HESCs. We employed targeted small interfering RNA (siRNA) to downregulate transcripts from the gene that encodes ERα, *ESR1*.

After treating HESCs with 20 nM of ESR1-targeted siRNA, we allowed the cells to differentiate for 72 hours under hypoxic conditions. We confirmed that *ESR1* transcripts were indeed reduced after targeted siRNA treatment compared to scrambled siRNA control (**Fig. 2A**) and found that there was a marked reduction of *HIF2α* transcripts (**Fig. 2B**), accompanied by a reduction in *RAB27B* gene expression (**Fig. 2C**). Finally, we collected the conditioned media from the treated cells, and via MRPS, demonstrated that *ESR1* siRNA treatment significantly reduced the number of EVs secreted from the differentiating HESCs (**Fig. 2D**). These data collectively illustrate that impairment of ERα signaling exhibits a gene expression and EV secretion trend similar to that of phthalate exposure.

**Fig. 2.**
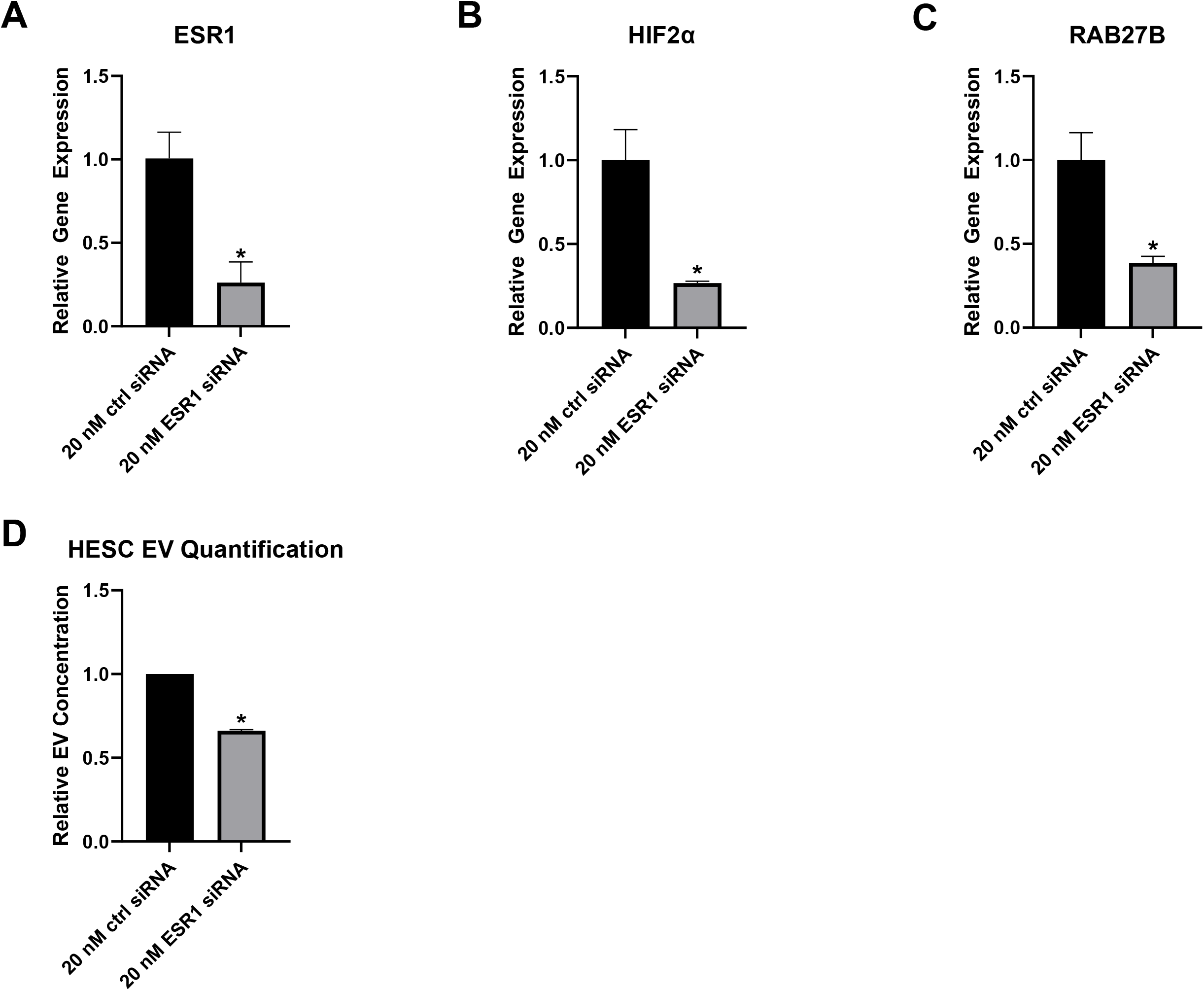
siRNA-mediated knockdown of *ESR1* in differentiating HESCs impairs EV secretion via the HIF2α/Rab27b pathway. HESCs were treated with 20 nM control or *ESR1*-specific siRNA for 24 hrs then grown in culture under hypoxic conditions with Decidualization Cocktail for 72 hrs. RNA was extracted and gene expression analysis was performed using primers specific for *ESR1* (**A**), *HIF2α* (**B**), or *RAB27B* (**C**). *36B4* was used to normalize gene expression. Data are shown as mean fold change ± SEM (n = 3-4 independent experiments), * p<0.05, relative to control siRNA treatment. **D.** EVs were isolated from conditioned media using the miRCURY (Qiagen) kit and analyzed by MRPS (Spectradyne, LLC). Concentration of EVs ranging from 60 to 400 nm was quantified. Data are shown as mean fold change ± SEM (n = 2 independent experiments), * p<0.05, relative to control siRNA treatment.

Due to the similarities in phenotype between the suppression of ERα signaling and phthalate exposure, we further clarified the mechanism by which phthalates disrupt the HIF2α/Rab27b/EV secretion pathway. ERα, like other steroid receptors, binds to specific response elements in the DNA, modulating the expression of nearby genes (34). Thus, we conducted ChIP-qPCR experiments to first confirm that ERα directly binds to DNA within the *HIF2α* gene regulatory region, and then determine if phthalate exposure potentially inhibits that interaction. We therefore examined ERα binding to several putative estrogen response elements (EREs) within the *HIF2α* regulatory region. In HESCs that were treated with DC plus vehicle for 72 hours under hypoxic conditions, we found that ERα exhibited strong binding to a site approximately 19.6 kb upstream of the *HIF2α* transcription start site (TSS) (**Fig. 3A**). Strikingly, upon addition of 1 μg/mL DEHP or MEHP to the DC, the DNA occupancy of ERα at the same putative ERE within the *HIF2α* regulatory region was completely ablated (**Fig. 3B-C**). These results suggested that DEHP or MEHP prevents ERα binding to the ERE site that controls estrogen-dependent expression of the *HIF2α* gene. Impairment in this gene expression leads to a concomitant reduction of downstream expression of the *RAB27B* gene and EV secretion.

**Fig. 3.**
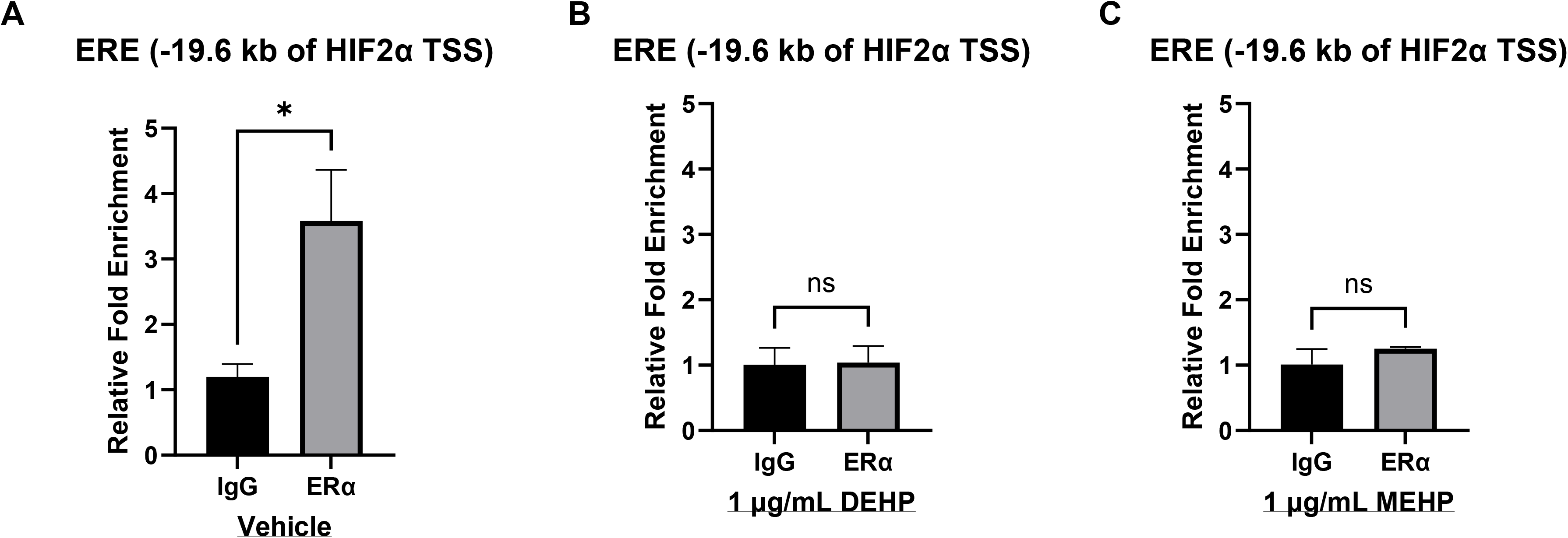
ERα binding to *HIF2α* promoter region in differentiating HESCs is impaired by treatment with DEHP or MEHP. ChIP was performed on HESCs that had been treated with Decidualization Cocktail and vehicle (**A**), 1 μg/mL DEHP (**B**), or 1 μg/mL MEHP (**C**) for 72 h under hypoxic conditions. Chromatin enrichment was quantified by qPCR and normalized to a negative control locus. Primers flanking putative ERE upstream of the *HIF2α* transcription start site were used to quantify chromatin enrichment. Data are shown as mean fold change ± SEM (n = 3 independent experiments), * p<0.05, relative to IgG control.

### DEHP or MEHP binds to ERα only exceedingly weakly

We next tested the possibility that the effects of DEHP or MEHP are mediated through direct binding of these phthalates to ERα, impairing its binding to the *HIF2α* regulatory region. Other EDCs have an affinity for ERα, such as bisphenol A (BPA) (weak affinity) or diethylstilbestrol (DES) (strong affinity), and can outcompete estradiol for binding to ERα at environmentally relevant concentrations (35). The binding of xenobiotic chemicals to ERα can inhibit receptor dimerization and the affinity of ERα for binding to DNA in a ligand-dependent manner (36). To test whether DEHP or MEHP can outcompete estradiol for receptor binding, we performed a competitive radiometric binding assay using purified full-length human ERα as described previously (29). As predicted, positive control of cold estradiol was able to outcompete tritiated estradiol with an IC50 of ∼0.9 nM (**Fig. 4A**), while DEHP and MEHP, with binding affinities about a million fold lower, were only able to outcompete tritiated estradiol at supraphysiological concentrations with an IC50 of ∼0.5 mM (**Fig. 4B-C**), much higher than the concentration at which we treated HESCs (1 μg/mL ≈ 2.5 μM DEHP or 2.9 μM MEHP). Additionally, if we had observed strong binding and interruption of activity of ERα by DEHP or MEHP, we would have expected a similar transcriptional response to phthalate exposure in all genes regulated by ERα in HESC. However, the expression of *PGR*, a known ERα-regulated gene, remained unaffected following phthalate exposure (**Fig. 1B**), indicating that the gene regulatory function of ERα is not controlled by direct binding of phthalates to this receptor.

**Fig. 4.**
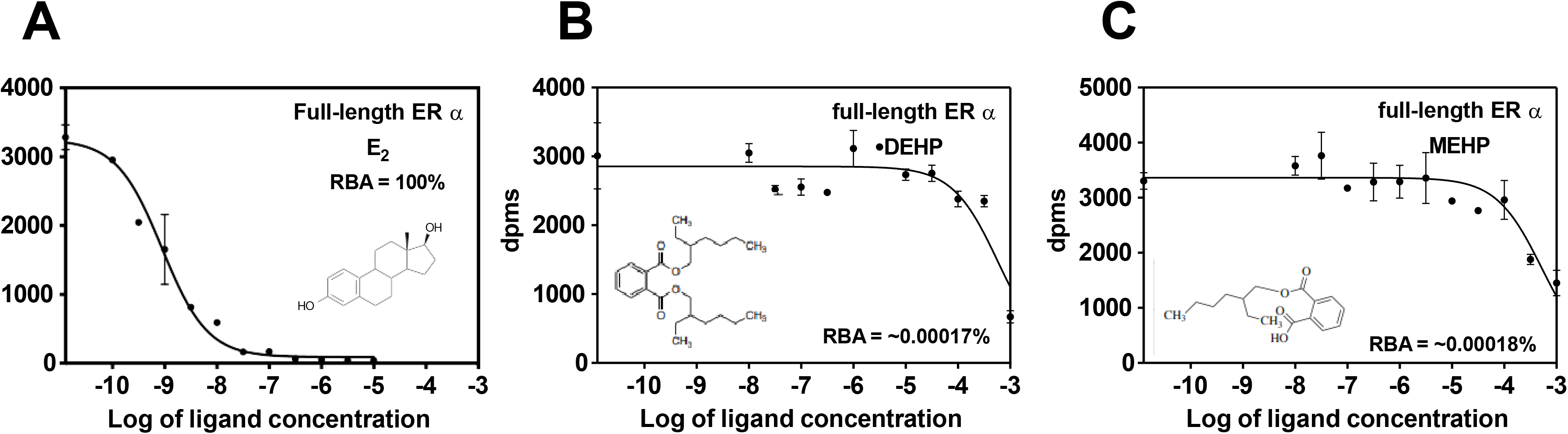
DEHP and MEHP are unable to efficiently outcompete estradiol for ERα occupancy. A radiometric competitive binding assay was performed using baculovirus-expressed human full-length ERα (Thermo Fisher Scientific) and tritiated estradiol. Relative binding affinities were quantified as increasing concentrations of competitor ligands estradiol (**A**), DEHP (**B**), or MEHP (**C**) were added to measure the amount of tritiated estradiol that was being displaced. Data are shown as mean dpm (disintegrations per minute) ± SEM (n = 3 replicates), and IC50 is shown. RBA (relative binding affinity) is calculated relative to estradiol, and the RBA for estradiol is defined as 100. The Kd for estradiol is 0.2 nM, and the Ki (in nM) for the competitors DHEP and MHEP can be determined from Ki = 0.2 x 100/RBA.

### Exposure to phthalates induces hypermethylation at CpG island in HIF2α gene regulatory region

Since DEHP has been associated with inducing differential DNA methylation (18,20,37–40), we investigated whether DEHP or MEHP changes the methylation status of DNA within the *HIF2α* regulatory region, preventing ERα from properly binding and regulating *HIF2α* expression in HESC. We tested this possibility by designing primers surrounding the putative ERE site where we had identified the impaired ERα binding (-19.6 kb from *HIF2α* TSS), as well as within two CpG Islands (-17.6 kb to −17.2 kb and −8.1 kb to −7.5 kb) located within the regulatory region as identified using a method previously described (30). A schematic detailing the location of these DNA regions is provided in **Fig. 5A**. We then used OneStep qMethyl-PCR (Zymo) to quantify the percent of methylated DNA within the regions spanned by our primers. We found that there was no change in DNA methylation surrounding the −19.6 kb putative ERE (**Fig. 5B**). Also, within the −17.6 kb CpG island, there appears to be no change DNA methylation, although MEHP may have a partial effect (**Fig. 5C**). Interestingly, within the −8.1 kb CpG island, we found that both DEHP and MEHP treatment induced a significant increase in DNA methylation compared to vehicle control (**Fig. 5D**).

**Fig. 5.**
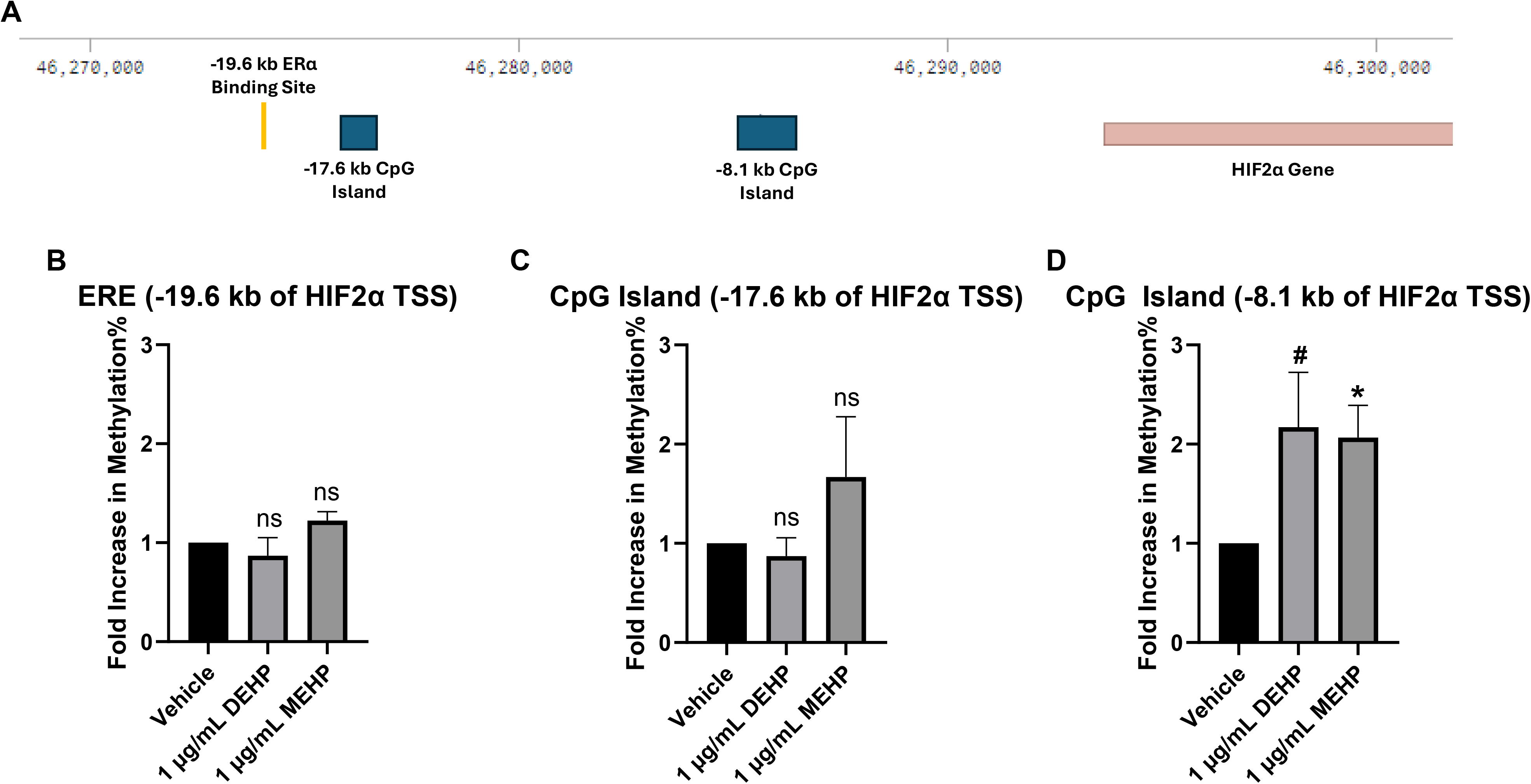
Treatment of differentiating HESCs with DEHP or MEHP alters DNA methylation patterns within *HIF2α* regulatory region. **A.** Schematic detailing *HIF2α* regulatory region. Browser window from Benchling.com. **B-D.** HESCs were treated with a Decidualization Cocktail and vehicle, 1 μg/mL DEHP, or 1 μg/mL MEHP for 72 h under hypoxic conditions. DNA was extracted and changes to the genomic DNA methylation status of CpG islands and EREs within the *HIF2α* regulatory region were quantified using OneStep qMethyl-PCR (Zymo). Data shown as mean fold change in methylation % compared to vehicle ± SEM (n = 2-3 independent experiments), * p<0.05, # p = 0.067 relative to vehicle control.

### Phthalate exposure disrupts SP1-mediated DNA binding of ERα in HIF2α gene regulatory region

To better understand how increased DNA methylation within the −8.1 kb CpG island disrupts ERα binding at the −19.6 kb putative ERE, we performed *in silico* analyses to determine what transcription factors were predicted to bind within the −8.1 kb CpG island. Notably, we discovered that specificity protein 1 (Sp1) was expected to bind within the CpG island at a site noted in our schematic (**Fig. 6A**). We therefore designed primers surrounding the putative Sp1 binding site. We conducted ChIP-qPCR to assess Sp1’s binding to its predicted site within the CpG island in differentiating HESCs. Indeed, we found that Sp1 showed significant DNA occupancy within the −8.1 kb CpG island (**Fig. 6B**). Most importantly, Sp1 was no longer able to occupy the DNA after exposure to DEHP or MEHP (**Fig. 6C-D**), likely due to the increased DNA methylation that we have exhibited occurs within that CpG island after phthalate exposure.

**Fig. 6.**
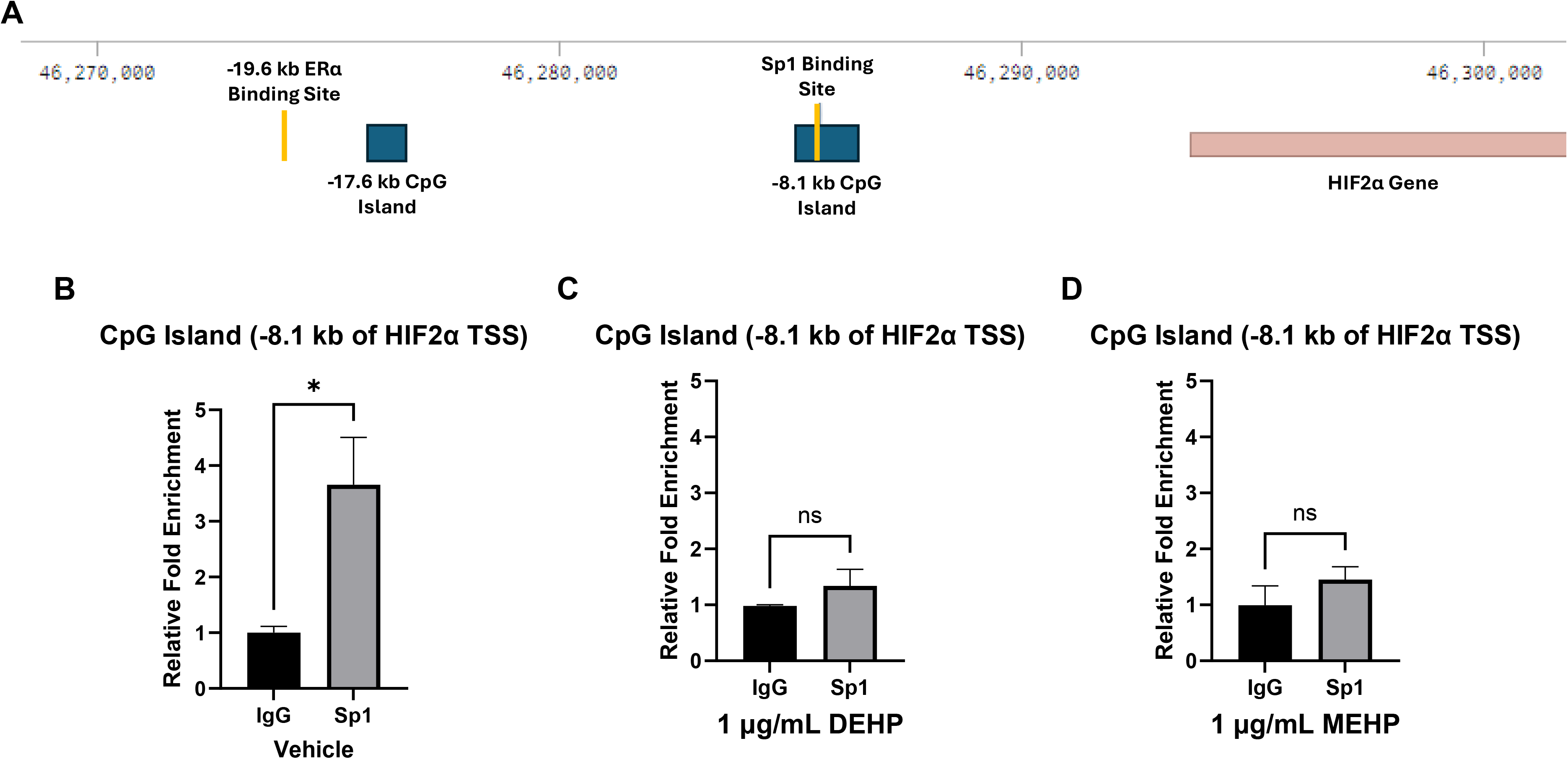
Sp1 binding to −8.1 kb CpG Island in differentiating HESCs is impaired by treatment with DEHP or MEHP. **A.** Schematic detailing *HIF2α* regulatory region and indicating Sp1 binding site. Browser window from Benchling.com. ChIP was performed on HESCs that had been treated with Decidualization Cocktail and vehicle (**B**), 1 μg/mL DEHP (**C**), or 1 μg/mL MEHP (**D**) for 72 h under hypoxic conditions. Chromatin enrichment was quantified by qPCR and normalized to a negative control locus. Primers flanking putative Sp1 binding sites within CpG island were used to quantify chromatin enrichment. Data are shown as mean fold change ± SEM (n = 3 independent experiments), * p<0.05, relative to IgG control.

Sp1 is well known for its interaction with ERα, modulating gene expression in coordination with ERα either by stabilizing ERα binding to a putative ERE or via a non-classical tethering interaction where ERα is not bound to DNA (41). Therefore, we hypothesized that, in this case, Sp1 binding at the −8.1 kb CpG island might stabilize ERα binding at the −19.6 kb putative ERE, an interaction made possible by DNA looping which is characteristic of ERα transcriptional regulation of many genes (42). It has been previously described that this DNA looping phenomenon occurs around the HIF2α gene (43). To test this hypothesis, we first determined if *HIF2α* gene expression is altered in the absence of Sp1. We downregulated *SP1* transcripts by siRNA-mediated knockdown (**Fig. 7A**) and exhibited that this knockdown led to a decrease in *HIF2α* gene expression (**Fig. 7B**). Further, via ChIP-qPCR, we showed that knockdown of *SP1* prohibited occupancy of ERα at the −19.6 kb putative ERE (**Fig. 7C-D**), supporting our hypothesis that ERα transcriptional regulation of HIF2α is reliant on Sp1-dependent stabilization of ERα DNA binding at a site 19.6 kb upstream of *HIF2α* TSS.

**Fig. 7.**
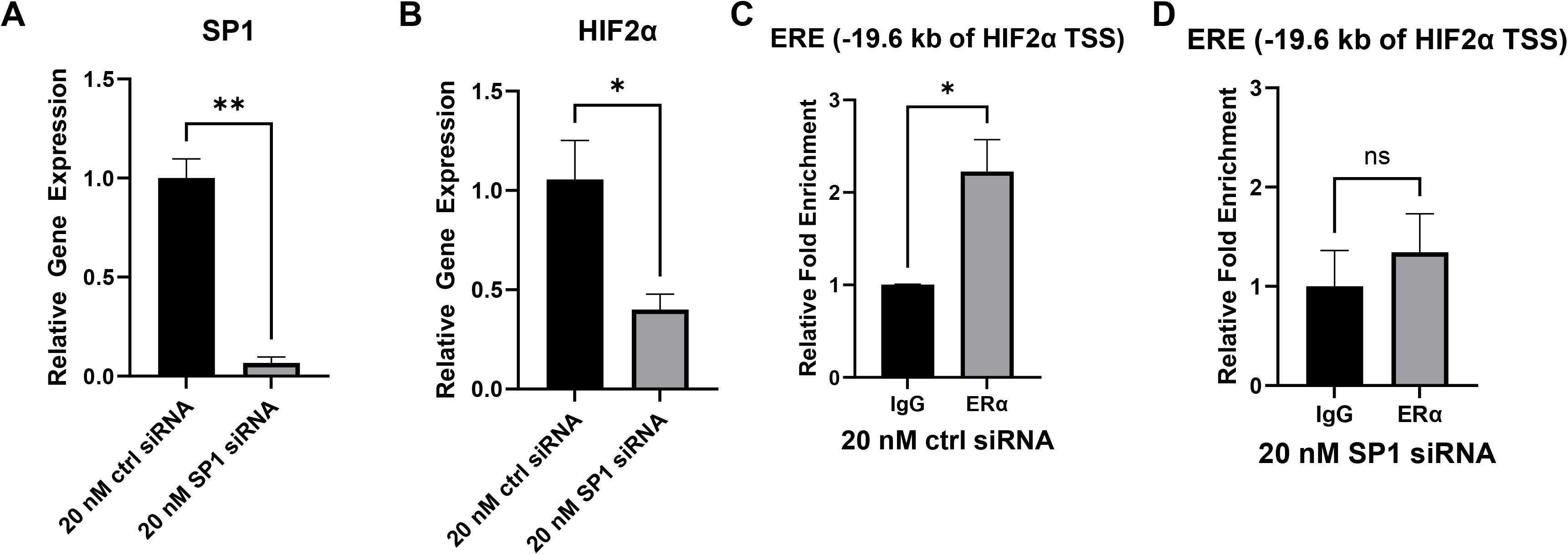
ERα binding to putative ERE and transcriptional regulation of the *HIF2α* gene is dependent upon Sp1. HESCs were treated with 20 nM control or *SP1*-specific siRNA for 24 h then grown in culture under hypoxic conditions with a Decidualization Cocktail for 72 h. RNA was extracted, and gene expression analysis was performed using primers specific for *SP1* (**A**) and *HIF2α* (**B**). *36B4* was used to normalize gene expression. Data are shown as mean fold change ± SEM (n = 2-3 independent experiments), * p<0.05, **p<0.01, relative to control siRNA treatment. ChIP was performed on HESCs that had been treated with Decidualization Cocktail for 72 h under hypoxic conditions and had previously been treated with 20 nM control (**C**) or *SP1*-specific (**D**) siRNA. Chromatin enrichment was quantified by qPCR and normalized to a negative control locus. Primers flanking putative ERE upstream of the *HIF2α* transcription start site were used to quantify chromatin enrichment. Data are shown as mean fold change ± SEM (n = 3-4 independent experiments), * p<0.05, relative to IgG control.

### DNA methyltransferase gene expression in HESC is altered by phthalate exposure

Finally, we identified a plausible mechanism by which phthalate exposure may induce hypermethylation at certain sites in the DNA of differentiating HESCs. While it has been well documented that phthalates can cause epigenetic alterations, including DNA methylation, as mentioned above (18,20,37–40), few studies offer insights into how phthalate exposure may alter DNA methylation. It is difficult to determine a universal mechanism describing how an environmental insult, such as phthalate exposure, can influence DNA methylation due to the cell- and tissue-specific nature of DNA methylation (44). However, methyl groups are conferred and maintained on the DNA in most cell types by a family of DNA methyltransferases (DNMTs) (45). Therefore, in differentiating HESCS, we performed qRT-PCR to determine if the expression of any DNMTs changed. Interestingly, we found that maintenance methyltransferase *DNMT1* gene expression was markedly enhanced after exposure to DEHP or MEHP (**Fig. 8A**), while *de novo* methyltransferases *DNMT3A* and *DNMT3B* gene expression was unaffected (**Fig. 8B-C**), raising the possibility that phthalate-induced upregulation of DNMT1 in HESC might be involved in driving the observed DNA hypermethylation.

**Fig. 8.**
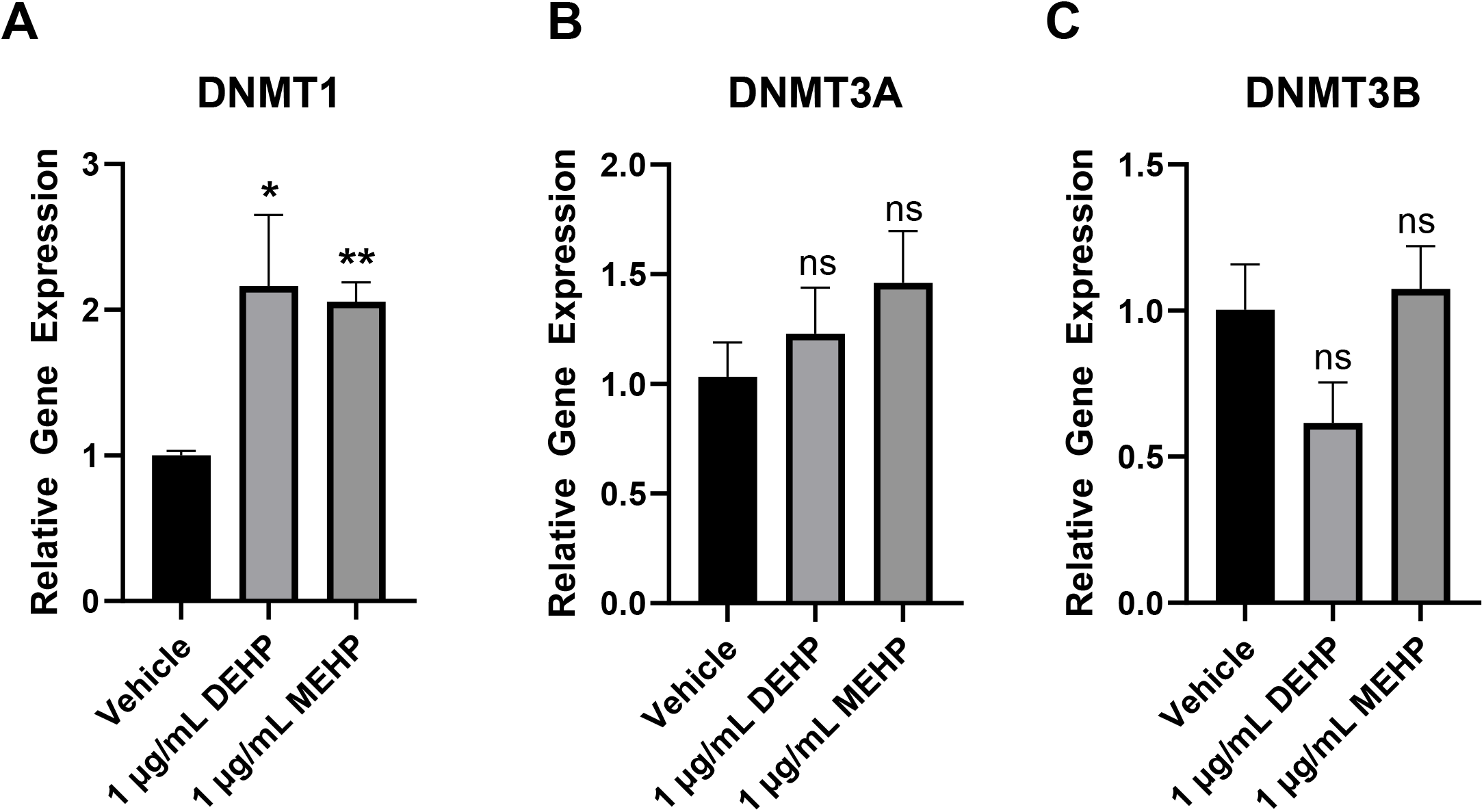
Phthalate exposure alters DNA methyltransferase gene expression. HESCs were treated with a Decidualization Cocktail and vehicle, 1 μg/mL DEHP, or 1 μg/mL MEHP for 72 hrs under hypoxic conditions. RNA was extracted and gene expression analysis was performed using primers specific for DNA methyltransferase genes, *DNMT1* (**A**), *DNMT3A* (**B**), and *DNMT3B* (**C**). *36B4* was used to normalize gene expression. Data are shown as mean fold change ± SEM (n = 3-6 independent experiments), * p<0.05, **p<0.01 relative to vehicle treatment.

## Discussion

Epidemiological studies associate DEHP with several pathophysiological conditions, including those of the female reproductive tract, leading to numerous countries placing restrictions on the use of DEHP in products such as toys and certain food contact materials (46). Despite these restrictive measures, DEHP remains ubiquitous in the environment with DEHP metabolites being found in the urine of nearly all tested individuals (2–4). Exposure to DEHP is a significant concern, particularly during the vulnerable period of pregnancy. In this study, we investigated the impact of DEHP exposure on disrupting a crucial cell-cell communication mechanism in the uterus during early pregnancy. Our analysis revealed that exposure to DEHP or its metabolite MEHP impairs the regulation of *HIF2α* gene expression in HESCs undergoing decidualization under hypoxic conditions, reducing EV secretion by these cells.

Our previous studies described how the downregulation of *HIF2α* led to decreased *RAB27B* expression, perinuclear accumulation of secretory granules, and a reduction in EV secretion (22,25,26). In the current study, we found that *RAB27B* expression decreased in differentiating HESCs treated with phthalates, resulting in a reduced number of EVs secreted into the conditioned medium. This finding is significant because EVs are believed to play a vital role in facilitating angiogenesis, trophoblast differentiation, and other processes significant for ensuring reproductive success. Disruption of endometrial cell-to-cell signaling has been shown to impair the development of the placenta, which may result in various pathophysiologies, including recurrent miscarriage (49–51), which is a condition that has also been linked to phthalate exposure (9). To our knowledge, this is the first report describing impairment of EV secretion by phthalate exposure.

One of our most striking findings was the demonstration that DEHP and MEHP treatment affects the estrogenic regulation of the HIF2α/Rab27b/EV secretion pathway in decidualizing HESCs.

Estrogen has been shown to induce *Hif2α* expression in mice (22). We report here that the knockdown of ERα resulted in the downregulation of *HIF2α* expression, resulting in reduced *RAB27B* expression and EV secretion. Given the similar patterns of gene expression and EV secretion observed after treatment with DEHP or MEHP, we tested the hypothesis that phthalate exposure may impair the regulation of HIF2α expression by ERα. Consistent with this hypothesis, we found that DEHP and MEHP exposure prohibits ERα from binding to an ERE sequence −19.6 kb upstream of the *HIF2α* gene. EREs are often found >5kb upstream of a gene, regulating expression via DNA looping (42,52,53,54). Interestingly it was previously reported that *HIF2α* is regulated by ERα via DNA looping interactions (43,55).

We considered the possibility that DEHP and MEHP may be binding to ERα, which would prevent estrogen from binding and inducing nuclear localization of ERα, consequently reducing ERα occupancy of DNA within the *HIF2α* regulatory region. Our radiometric estradiol competition binding assay confirmed that only at supraphysiological conditions could DEHP or MEHP outcompete estradiol for ERα occupancy. These data are supported by previous reports that only at non-environmentally relevant concentrations of DEHP can outcompete estradiol for ERα binding (56). Additionally, *in vitro* reporter gene assays had described DEHP to display no estrogenic activity (57,58). Therefore, we concluded that phthalates inhibit estrogenic regulation of *HIF2α* not by directly binding to ERα but via another mechanism.

Exposure to DEHP has been shown to induce changes in DNA methylation in several studies (18,20,37–40). The *HIF2α* gene has been shown to have varying DNA methylation patterns, with different populations displaying altered methylation depending on their duration of residence at high altitudes (59). Often, changes to DNA methylation can be seen only after long-term exposure to an insult. However, hypoxia has been shown to drastically speed up the hypermethylation process with measurable changes to the methylation state and expression of genes appearing within mere hours (60). Taking this into consideration, we employed a qMethyl- PCR kit (Zymo) to measure the percentage of methylated DNA in regions spanned by our primers. We demonstrated that, although no DNA methylation changes were induced at the -20kb ERE site, the CpG island at −8.1 kb showed increased DNA methylation in response to both DEHP and MEHP.

We considered the possibility that DNA hypermethylation in the −8.1 kb CpG island could be responsible for prohibiting ERα binding to the −19.6 kb ERE. We found DNA motifs within the CpG island that predicted that the transcription factor Sp1 could bind there. We confirmed Sp1 occupancy of the DNA within the CpG island and then showed that it is ablated by exposure to DEHP or MEHP. Further, we showed, using siRNA-mediated knockdown of *SP1*, that ERα DNA occupancy at −19.6 kb and subsequent regulation of *HIF2α* gene expression is dependent upon the presence of Sp1 at its binding site. The collaboration between Sp1 and ERα in gene expression transactivation is well-documented (52,54,64). We, therefore, posit that the binding of Sp1 to the −8.1 kb GC-rich DNA motif allows for interaction with ERα, stabilizing its binding to the ERE. In this instance, an interaction between Sp1 and ERα would require DNA looping, which is quite common in ERα-regulated genes (52,54,65).

These data provide the basis of the mechanism suggested by this study. That is, phthalate exposure induces abnormal DNA methylation in a CpG island −8.1 kb of the *HIF2α* gene, inhibiting transcription factor Sp1 from binding to its consensus sequence located within the CpG island. The lack of Sp1 binding at the CpG island prevents Sp1 from facilitating the binding of ERα to its binding site −19.6 kb of the *HIF2α* gene, subsequently preventing ERα from activating *HIF2α* gene expression. The resultant decreased HIF2α expression then reduces the gene expression of its downstream target, vesicular trafficking protein RAB27b, thereby leading to diminished EV secretion into the extracellular space (**Fig. 9**).

**Fig. 9.**
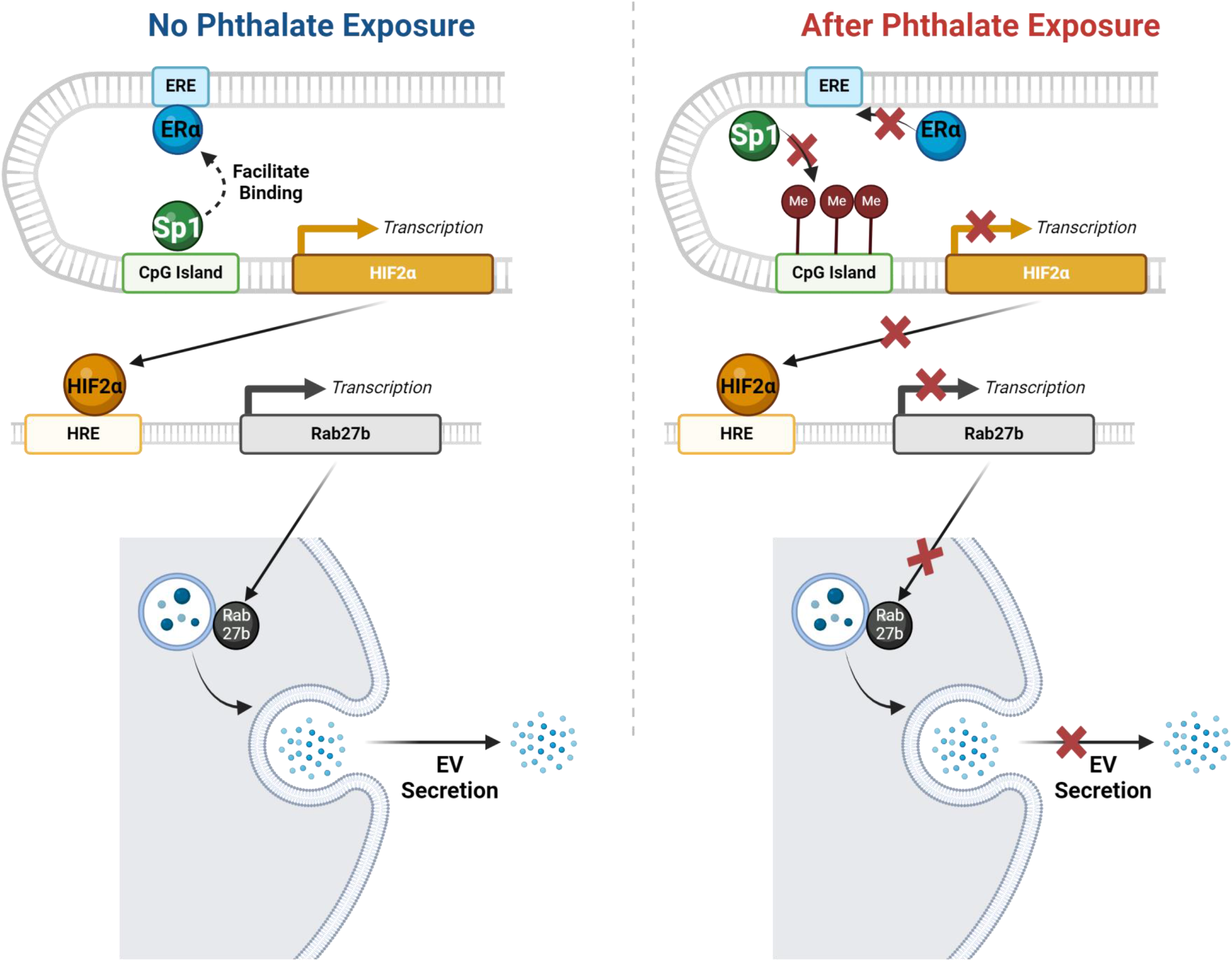
Summary of the proposed mechanism of phthalate-induced disruption of estrogenic regulation of EV signaling in differentiating HESCs. Left panel illustrates estrogenic regulation of EV secretion without exposure to phthalate. Sp1 binds to CpG island in *HIF2α* regulatory region, facilitating ERα binding to its ERE. This leads to enhanced transcription of *HIF2α* gene. HIF2α protein then regulates the expression of *RAB27B* gene, and RAB27B protein traffics the multivesicular body to the cell membrane, allowing for the release of EVs. Right panel illustrates disrupted estrogenic regulation of EV secretion after exposure to phthalate. Increased methylation of DNA in CpG island in *HIF2α* regulatory region prevents Sp1 binding, subsequently preventing Sp1 from facilitating ERα binding. This leads to a decrease in expression of *HIF2α* and downstream gene *RAB27b*, ultimately leading to decreased EV secretion.

Epigenetic regulation of gene expression by DNA methylation is a complex process, involving *de novo* and maintenance of DNA methyltransferases, methyl donors, demethylases, and methyl-CpG binding proteins. Abnormal expression or activity of any of these factors could lead to the observed changes in DNA methylation after exposure to phthalates. It has been suggested that, in ER+ breast cancer cells, exposure to DEHP can cause an increase in *DNMT1* gene expression, which ultimately leads to DNA hypermethylation (66). Our data showed a similar response by HESCs, with *DNMT1* expression being significantly enhanced in the presence of DEHP or MEHP. DNMT1 is typically recognized as a maintenance methyltransferase, which primarily adds methyl groups to hemimethylated DNA. However, it also has a weaker affinity for unmethylated DNA, enabling it to perform *de novo* methylation. (67). It is plausible that, due to increased *DNMT1* expression after phthalate exposure, DNMT1 is responsible for the increased DNA methylation we have observed in the −8.1 kb CpG island. However, we realize that the mechanism of action linking phthalate exposure and DNA methylation is likely to be more complex, involving additional activities such as demethylases.

In summary, we provide a relevant example of how phthalates disrupt gene expression in HESC, leading to impairment in EV secretion and cell-to-cell communication. Furthermore, we anticipate that exposure to phthalates also triggers genome-wide alterations in DNA methylation. Our findings lay a solid foundation for future studies aimed at investigating how phthalate- induced changes in DNA methylation may affect the expression of other critical genes active in the uterus during early pregnancy. Many of these genes are similarly regulated by the coordination of hormone receptors and other cofactors. Continued analysis of phthalate-induced alterations in the genomic and epigenomic landscapes of differentiating endometrial stromal cells—essential for establishing and maintaining pregnancy—would provide deeper insights into the reasons why phthalates have been linked to adverse pregnancy outcomes.

